# Elucidating dynamics and regulation of alternative splicing in osteogenic differentiation

**DOI:** 10.1101/2020.10.30.362384

**Authors:** Yuanyuan Wang, Rene F Chun, Samir Adhikari, Christopher M Lopez, Mason Henrich, Vahe Yacoubian, Lan Lin, John S. Adams, Yi Xing

**Affiliations:** Bioinformatics Interdepartmental Graduate Program, University of California, Los Angeles; Department of Orthopaedic Surgery, University of California, Los Angeles; Department of Molecular, Cell & Developmental Biology, University of California, Los Angeles; Center for Computational and Genomic Medicine, Children’s Hospital of Philadelphia; Department of Pathology and Laboratory Medicine, Perelman School of Medicine, University of Pennsylvania

## Abstract

Nearly all human multi-exonic genes undergo alternative splicing (AS) via regulation by RNA-binding proteins (RBPs), but few studies have examined the temporal dynamics of AS and its regulation during cell differentiation in the bone niche. We sought to evaluate how AS, under the control of RBPs, affects cell fate commitment during induced osteogenic differentiation of human bone marrow-derived multipotent stem/stromal progenitor cells (MSPCs). We generated a time-course RNA sequencing (RNA-seq) dataset representative of induced MSPC differentiation to osteoblasts. Our analysis revealed widespread AS changes, coordinated with differential RBP expression, at multiple time points, including many AS changes in non-differentially expressed genes. We also developed a computational approach to profile the dynamics and regulation of AS by RBPs using time-course RNA-seq data, by combining temporal patterns of exon skipping and RBP expression with RBP binding sites in the vicinity of regulated exons. In total we identified nine RBPs as potential key splicing regulators during MSPC osteogenic differentiation. Perturbation of one candidate, *KHDRBS3*, inhibited osteogenesis and bone formation *in vitro*, validating our computational prediction of “driver” RBPs. Overall, our work highlights a high degree of complexity in the splicing regulation of MSPC osteogenic differentiation. Our computational approach may be applied to other time-course data to explore dynamic AS changes and associated regulatory mechanisms in other biological processes or disease trajectories.

## Introduction

Alternative splicing (AS) is a prevalent mechanism for generating regulatory and functional complexity in eukaryotes (Nilsen and Graveley, 2010). The most common mode of AS in human cells is exon skipping. Less common modes include use of alternative 5’ splice sites, alternative 3’ splice sites, mutually exclusive exons, and intron retention (McManus and Graveley, 2011; Park et al., 2018). In each case, the resultant transcript isoforms might produce proteins with altered structure and function. Not only does AS have a vital role in normal cellular processes (Cieply et al., 2016; Song et al., 2017; Zhang et al., 2016), it is implicated in a variety of diseases (Dvinge et al., 2016; Pagani and Baralle, 2004; Rahman et al., 2020; Scotti and Swanson, 2016; Zhang and Manley, 2013).

Splicing is coordinately regulated by the interaction of *cis*-acting RNA elements and *trans*-acting splicing factors. The cis-acting elements include pre-mRNA sequence motifs in exons or flanking introns that function as splicing enhancers or silencers when bound by RNA-binding proteins (RBPs) acting as splicing factors (Gerstberger et al., 2014). Recently, the study of AS has been accelerated by the development of high-throughput sequencing methods and novel bioinformatic tools that identify and quantitate AS (Park et al., 2018; Shen et al., 2014).

As humans age the balance of bone and fat content in the skeleton shifts more toward fat, leading the way to skeletal fragility (Duque, 2008; Meunier et al., 1971). Age-related bone loss resulting in osteoporosis and increased fractures represents a major health problem with increased morbidity and decreased quality of life (Blume and Curtis, 2011; Kapinos et al., 2018). Human bone-forming cells or osteoblasts are derived from bone marrow-derived multipotent stem/stromal progenitor cells (MSPCs). As their name implies, MSPCs are capable of undergoing osteogenesis, adipogenesis, chondrogenesis and myogenesis (Augello et al., 2010; Pittenger et al., 1999) with a major differentiation branch point between osteogenesis and adipogenesis. This branch point represents a binary fate choice for MSPC. The signals that specify MSPC to specific cell differentiation pathways are not fully understood but transcription factors are considered by most to be the dominant signal. *RUNX2* (Komori et al., 1997; Otto et al., 1997) and *PPARγ* (Barak et al., 1999; Rosen et al., 1999) are well known transcription factors that have roles in osteogenesis and adipogenesis, respectively. Our understanding of osteogenic differentiation from MSPC has made tremendous progress in the last several decades. This includes: 1) establishing criteria for MSPC characterization (Chan et al., 2018; Dominici et al., 2006; Robey and Riminucci, 2020); 2) optimizing procedures for isolation (Soleimani and Nadri, 2009; Wagey and Short, 2013) and culture of MSPC (Ciuffreda et al., 2016; Jaiswal et al., 1997); and 3) determining some of the key regulatory molecules, particularly those that control transcription, that modulate MSPC differentiation patterns (Chen et al., 2016; Han et al., 2019).

In the mouse, recent studies in transcription and epigenetic mechanisms regulating MSPC differentiation using single-cell RNA sequencing (scRNA-seq) approaches have yielded important findings with regard to the fate decisions in MSPC differentiation (Rauch et al., 2019; Wolock et al., 2019; Zhong et al., 2020). For example, Zhong et al identified a lineage commitment progenitor population that could differentiate to adipocytes or osteocytes (Zhong et al., 2020). Additionally, two groups (Rauch et al., 2019; Wolock et al., 2019) found that the expression profiles of transcription factors revealed that adipogenesis required a much larger number of expressed transcription factors compared to osteogenesis, and that there existed a subset of transcription factors that acted in both pro-osteogenic and anti-adipogenic manner, supporting a net gain in osteoblasts over adipocytes. One report demonstrated that adipogenesis invoked more chromatin remodeling relative to osteogenesis (Rauch et al., 2019). In each report noted above, selection of either the osteogenesis or adipogenesis differentiation path appeared binary.

In this work, we studied the role of AS during the MSPC osteogenic differentiation. We conducted a time course study with primary MSPCs derived from the marrow space of human femurs cultured in osteogenic media. At selected time points, RNA was obtained for RNA sequencing (RNA-seq). Extensive and dynamic transcriptomic alterations are orchestrated during induced osteogenic differentiation, including remodeling of the gene expression and splicing programs. We developed a multi-step bioinformatic strategy to identify RBPs that may be key regulators of AS in the process of osteogenesis.

## Results

### Extensive transcriptomic alterations characterize the induced, stepwise differentiation of primary MSPCs to osteoblasts

In order to evaluate gene expression and AS regulation during osteogenesis, we cultured MSPCs in osteogenic differentiation media over 12 days to obtain temporal MSPC osteogenic differentiation datasets. Cells were stained for alkaline phosphatase activity and mineralization by alizarin red every two days (**Figure 1A**). The anticipated increase in staining intensity was observed qualitatively (**Figure 1A**) and quantitatively (**Figure 1B**). RNAs were isolated in triplicate on day 0, day 2, day 4, day 6, day 8 and day 12, followed by high-throughput RNA-seq (**Figure S1A**). The RNA-seq-derived gene expression data showed that markers of immature osteoblasts (*MYC* and *SOX9*) were gradually downregulated as markers of osteoblast differentiation (*RUNX2* and *ALPL*) were upregulated during the time course (**Figure 1C**), consistent with osteogenic differentiation of the MSPCs. These sequential changes in histological staining and RNA-seq-derived marker gene expression confirmed MSPC progression toward a mature osteoblast.

**Figure 1.**
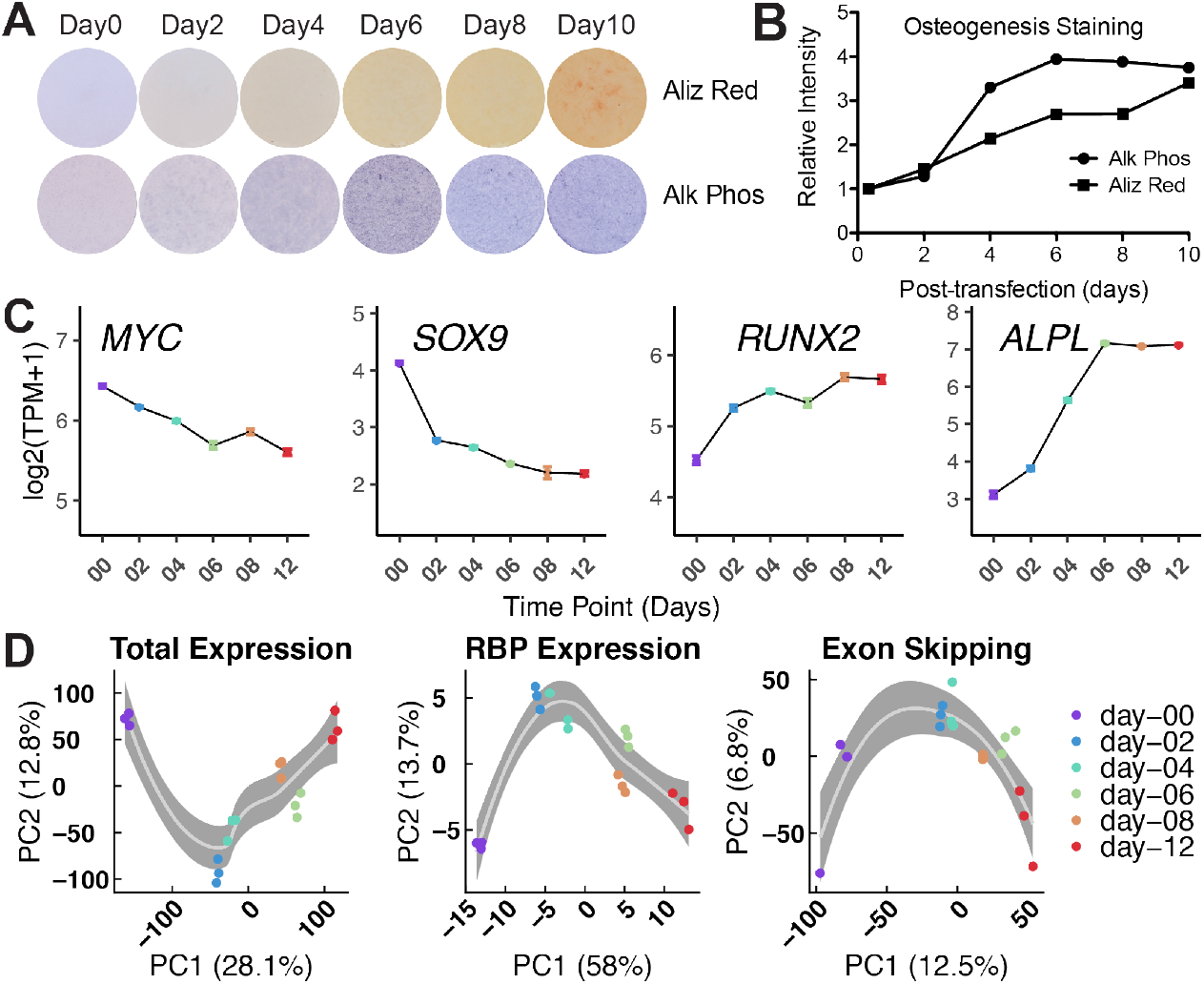
Extensive transcriptomic alterations characterize the induced stepwise differentiation of MSPC to osteoblasts. (A) Photographs of blue alkaline phosphatase and red hydroxyapatite (bone calcification) staining of MSPCs cultured in osteoblast differentiation medium over 10 days. (B) Measurement of optical density in photographed wells in panel A. (C) From-left-to-right are normalized expression measures from RNA-seq at indicated time points, showing a temporal decrease in immature marker (*MYC* and *SOX9)* expression and increase in osteoblast marker (*RUNX2* and *ALPL)* expression. Error bars represent the MEAN±SEM (n=3). (D) From left-to-right, principal component analysis (PCA) plots of total gene expression (**left**), RBP gene expression (**middle**), and exon skipping (**right**). Samples were projected to the space of the first two principal components (PCs) with percentage of variation explained shown in x- and y-axis labels. Local regression lines are added by LOESS (locally estimated scatterplot smoothing) method, with 95% confidence intervals shown in grey.

A substantial remodeling of the gene expression and AS profiles occurred in osteogenic media-induced osteogenic differentiation of MSPCs. Using rMATS-turbo (rMATS 4.0.2) (Shen et al., 2014), we uncovered ~23,000 AS events across all 18 samples (**Figure S1D**), including exon skipping, alternative 5’ splice sites, alternative 3’ splice sites and intron retention. Because exon skipping is the most prevalent and most well-characterized type of AS events in human transcriptomes (Park et al., 2018) and because it represented 72% of the total AS events in this study, we focused on exon skipping events (**Figure S1D**). Temporal profiling of induced changes in the MSPC transcriptome from RNA-seq was assessed globally by principal component analysis (PCA). The PCA plots of total gene expression, RBP gene expression, and exon skipping showed similar temporal patterns (**Figure 1D**), suggesting coordination between gene expression, RBP gene expression and exon skipping. Moreover, the genes contributing most to the separation of samples on PC1 in total gene expression PCA plot were highly enriched in RNA splicing-related gene ontology terms (**Figure S1B**, **S1C**), further indicating the coupling of gene expression and splicing programs during induced MSPC osteogenic differentiation. Interestingly, genes contributing most to the separation of samples on PC2 in exon skipping PCA plot were enriched in regulation of transmembrane calcium ion transport term as well as regulation of RNA splicing term (**Figure S1E**, **S1F**). This is consistent with the observation that the net flux of calcium to the extracellular environment in osteoblasts is closely related to mineralization of the collagenous extracellular matrix (Boonrungsiman et al., 2012), again highlighting the importance of AS in osteogenic differentiation. Together, this transcriptome-wide analyses of osteogenic differentiation identifies interplay of gene expression, splicing and bone development related biological processes.

### Pair-wise differential analysis identifies temporal patterns of gene expression and exon skipping during MSPC-to-osteoblast differentiation

It has been well established that coordinated splicing networks play a vital role in cell fate determination and result in physiological consequences in various developmental and tissue remodeling processes in humans (Baralle and Giudice, 2017). In the field of skeletal modeling and remodeling the focus has been on changes in transcription factor gene expression (Rauch et al., 2019). On the other hand, there is a lack of systematic assessment of splicing networks in skeletal development and maintenance. To fill this knowledge gap and elucidate the dynamics and regulation of AS during osteogenic differentiation, we performed pair-wise comparisons on both gene expression and exon skipping in MSPC over 12 days of induced osteogenesis (**Figure 2A**). Beginning in the early stages of osteogenesis (day 0-2), there was a robust and dynamic alteration in the splicing program of MSPCs as they matured to an osteoblast phenotype (**Figure 2A**).

**Figure 2.**
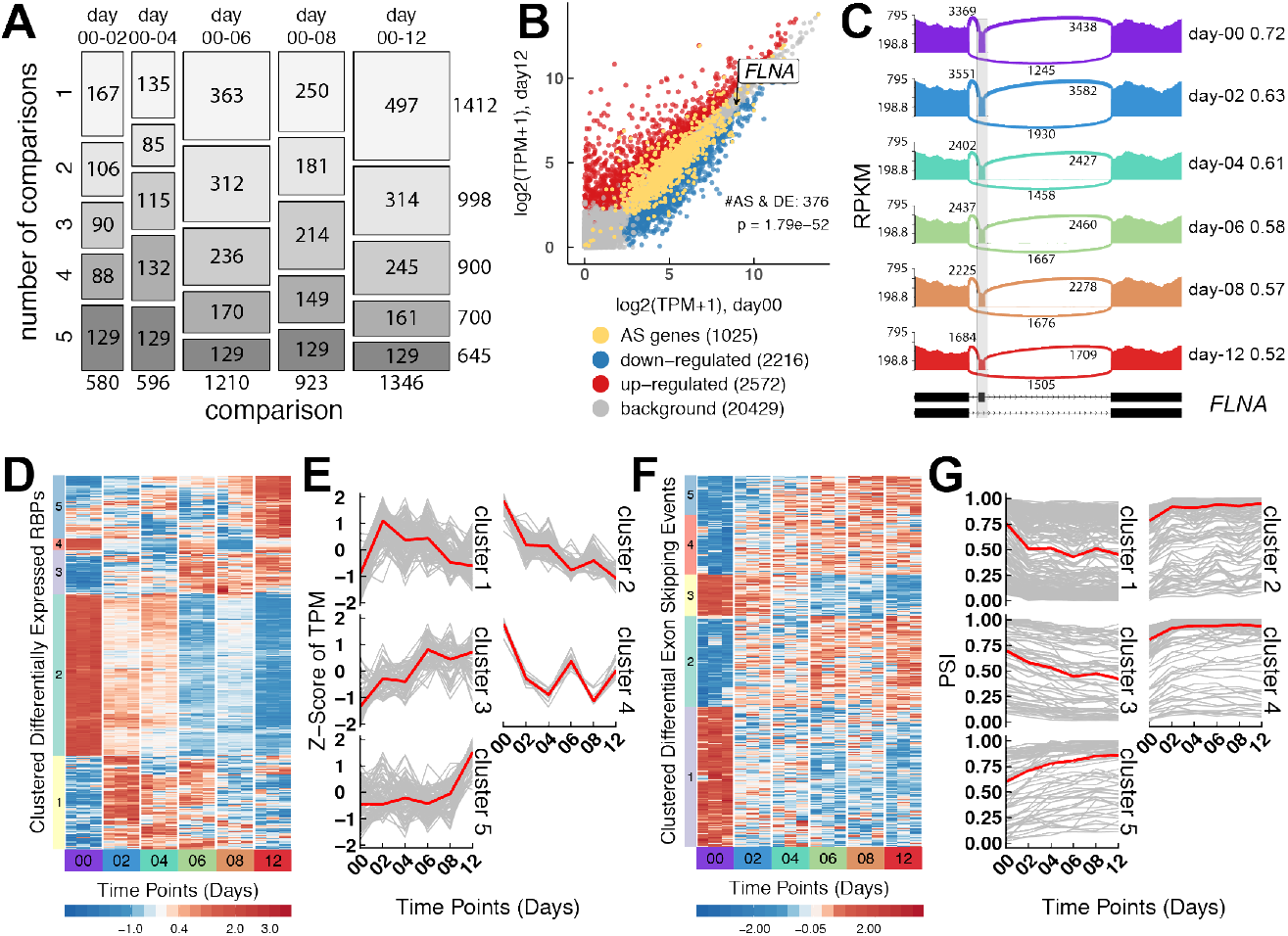
Pair-wise differential analysis identifies temporal patterns of gene expression and exon skipping during MSPC-to-osteoblast differentiation. (A) Mosaic plot display of significant exon skipping events in pair-wise comparisons with day 0. Each box is constructed by two factors, the specific comparison (x-axis) and total number of comparisons (y-axis) where events were identified as significant. The size of the box and number shown for each box represent the number of exon skipping events in each category. (B) Scatter plot comparing gene expression on day 0 vs. day 12. Red and blue dots represent up-regulated and down-regulated genes, respectively. Genes with significant exon skipping changes (denoted as AS genes) are depicted in yellow. Background genes, e.g., those with no differential gene expression and no exon skipping changes, are shown in grey. (C) Sashimi plot of *FLNA* gene from panel B is an example of a gene whose expression is not significantly changed but does harbor a significant change in exon skipping over time. The black bars and dashed lines on the bottom represent exons and introns, respectively. Solid peaks represent reads per kilobase per million mapped (RPKM) mapped to each region. Arches represent splice junctions and the numbers represents number of reads mapped to each splice junction. PSI values are indicated on the right side of the plot for each time point. (D-G) Heatmaps and line graphs illustrate the temporal coordination among RBP gene expression and exon skipping during induced osteogenic differentiation. Panel D, heatmap showing Z-score transformed transcripts per million (TPM) for 523 differentially expressed RBP genes; panel E, line plots revealing the patterns of change for corresponding clusters in panel D; panel F, heatmap showing Z-score transformed PSI values for 604 significantly changed exon skipping events; panel G, line plots revealing the patterns of change in PSI value for corresponding clusters in panel F. Red line represents the median value for each cluster.

To further decipher the relationship between gene expression and exon skipping in the process of induced MSPC osteogenic differentiation, we investigated the expression patterns of splicing-regulated genes.

Interestingly, most of those splicing-regulated genes were not differentially expressed (**Figure 2B**), suggesting that splicing, in and of itself, represents another layer of transcriptome remodeling during osteogenesis. For example, the filamin A gene (*FLNA*), of which missense point mutations are associated with a range of X-linked skeletal dysplasias (Feng and Walsh, 2004), was persistently highly expressed in both the MSPC and mature osteoblast population of cells with no change in gene expression over the 12-days of induced osteogenesis (**Figure 2B, 2C**). Despite no change in expression of *FLNA*, the inclusion level (percent spliced in, PSI) of the alternatively spliced exon 30 (ENST00000369850), which maps to the functional filamin domain repeat 15, steadily decreased from 0.72 to 0.52 over 12 days of osteogenic induction (**Figure 2C**).

Notably, of 604 exon skipping events (from 488 genes) with significant PSI value changes in at least 3 pair-wise comparisons, 53 were located in 47 genes encoding transcription factors (**Supplementary Table 1**). Those exon skipping-regulated transcription factors were often affected by 1) loss of a portion or the entirety of a functional domain encoded by the alternatively spliced exon, 2) a frame shift resulting in disruption or presence of downstream functional domains of the translated protein or 3) nonsense-mediated decay (**Supplementary Table 1**); either of these scenarios can lead to loss/gain of transcription factor function with a corresponding global reconstruction of the gene expression network. Exon skipping events residing in transcription factor genes without frameshift, NMD induction or functional domain encoding exons can also exert functional consequences. An example is differential inclusion/exclusion of exon 4 in the transcriptional co-activator gene, paired mesoderm homeobox protein 1 gene (*PRRX1*; also known as *PRX1*) (**Figure S2**). Two isoforms were produced by alternative splicing of the cassette exon 4 in *PRRX1*: the exon 4 skipping isoform, PRRX1A and exon 4 inclusion isoform, PRRX1B. Interestingly, the inclusion of exon 4 in the PRRX1 gene introduced an earlier stop codon, shifting exon 5 to the 3’ untranslated region and encodes a shorter protein isoform (PRRX1b) lacking the functional OAR (otp, aristaless, and rax) domain (**Figure S2A, S2B**). The OAR domain is involved in DNA binding and inhibition of transcription activation (Norris and Kern, 2001; Reichert et al., 2013). Previous studies have established that PRRX1a and PRRX1b act differentially to regulate progenitor cell proliferation and differentiation (Takano et al., 2016; Wang et al., 2018). Indeed, the sashimi plot in **Figure S2C** shows that the inclusion level of exon 4 decreased from 63% on day 0 to 41% on day 12, resulting in an isoform switch from the OAR-absent PRRX1B to the OAR-containing PRRX1A during induced MSPC osteogenesis. This suggests that OAR-containing PRRX1A favors an osteogenic differentiation fate more than does the OAR-absent PRRX1B. This result parallels with previous observations in the mouse system where overexpression of *Prrx1b*, but not *Prxx1a*, interferes with *Osx-* and *Runx2-*directed mRNA expression and inhibits pre-osteoblast-to-osteoblast differentiation (Lu et al., 2011).

Many AS events are coordinately regulated by *trans*-acting RBPs in a developmental stage-specific manner (Erhard et al., 2019; Fu and Ares, 2014). The global changes in RBP gene expression (**Figure 2D**) and exon skipping PSI values (**Figure 2F**) are displayed as heatmaps, each defined by 5 clusters with distinct inter-cluster temporal patterns determined by unsupervised hierarchical clustering. Overall, the most pronounced changes for both RBP gene expression and exon skipping were observed in the early stages of induced osteogenesis (from day 0 to day 2, **Figure 2D, 2F**). The temporal RBP gene expression patterns and temporal exon skipping PSI value patterns were further delineated into lines (**Figure 2E, 2G**). Distinct patterns of RBP gene expression emerged from different clusters (**Figure 2E**): steady upregulation (e.g., cluster 3); steady downregulation (e.g., cluster 2); early stage change (e.g., cluster 1) or late stage change (e.g., cluster 5). Distinct patterns of exon skipping PSI values were also observed (**Figure 2G**). This included clusters of exons with: progressive inclusion (e.g., cluster 5); progressive exclusion (e.g., clusters 3) and early stage change (e.g., clusters 1 and 2).

### Computational screening identifies RBP candidates for regulation of exon skipping during osteogenic differentiation

To further identify RBPs that could control the splicing program in induced MSPC osteogenic differentiation, we developed a computational screening method searching for potential ‘key splicing regulators’ among 129 RBPs with known motif position weight matrix (Dominguez et al., 2018; Ray et al., 2013). *Trans*-acting regulatory RBPs usually bind to *cis*-acting RNA elements in the precursor mRNA (pre-mRNA) in a sequence-specific manner with the resulting RBP-directed splicing behavior frequently dependent on the location of the RBP binding site relative to the regulated exon (Fu and Ares, 2014; Gabut et al., 2008). Taking this *cis-trans* locational information into consideration, the method described here combines both correlation analysis and region-specific motif enrichment analysis to estimate the region-specific regulatory potential of RBPs to influence pre-mRNA splicing (**Figure 3A**). Detected exon skipping events not significantly changed in any of the 5 pair-wise comparisons with day 0 were classified as background events, while exon skipping events that were significantly changed in at least three of the five comparisons were retrieved as foreground events. For each differentially expressed RBP, foreground events were assigned into a ‘positive’ or ‘negative’ foreground event set based on the correlation coefficient between its PSI value and the RBP gene expression if highly correlated (r^2^>0.5) (**Figure 3A, top**). Sequences extracted from each exon skipping event were subsequently scanned for the presence of an RBP binding site based on the motif scores calculated from matching the position weight matrix of the RBP motifs to possible binding positions (**Figure 3A, middle**). Finally, one-tailed Fisher’s exact tests were performed to estimate the region-specific enrichment of motifs in the foreground event sets compared to the background event set (**Figure 3A, bottom**). Eleven RNA binding motifs for nine RBPs were determined to be significantly enriched in at least one region for at least one foreground event set (**Figure 3B, 3C**).

**Figure 3.**
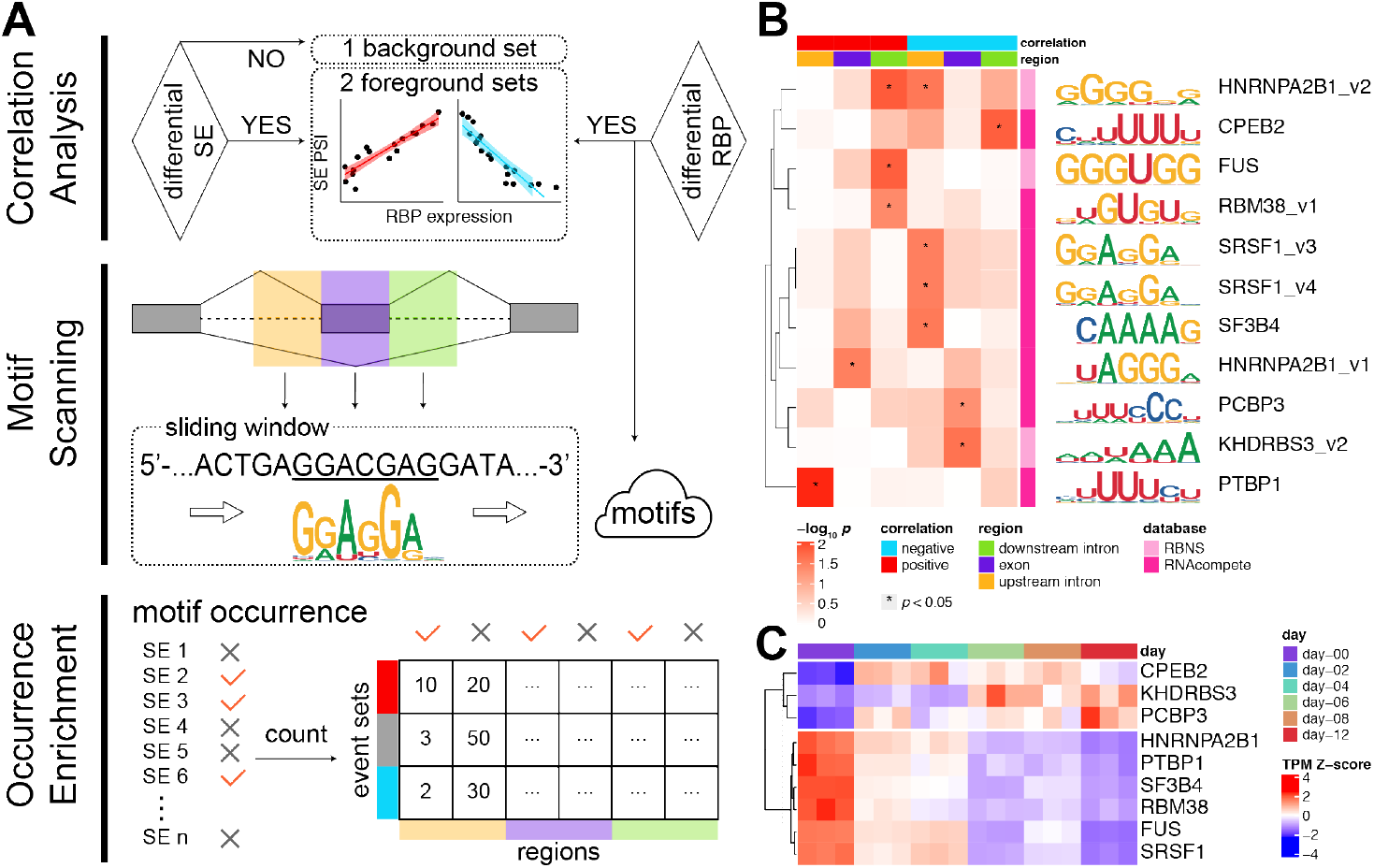
Computational screening identifies nine RBP candidate that may regulate exon skipping in osteogenic differentiation. (A) Schematic diagram describing the computational workflow for key splicing regulator screening. As depicted in the *top panel*, differential exon skipping analysis was performed to determine a background event set. For each differentially expressed RBP, significantly changing exon skipping events during osteogenesis were further split into two foreground sets, whose PSI change was positively (red, r^2^ > 0.5) or negatively (blue, r^2^ > 0.5) correlated with the corresponding change in RBP gene expression. As shown in the *middle panel*, a sliding window scanning tool was used to detect putative RBP RNA binding motifs in three regions in and around the skipped exon: 300 nt into the upstream intron (orange); the exon body (purple); and 300 nt into the downstream intron (green). The *bottom panel* shows a tabular registration of detected motif occurrence. One-tailed Fisher’s exact test was performed to ascertain the significance of motif occurrence enrichment in one of the three regions (upstream intron, exon body, or downstream intron) on the positive (red) or the negative (blue) foreground compared to background (grey). SE, exon skipping. (B) Heatmap depicting the log-transformed *p* values of 11 candidate RBP motifs (y-axis) exhibiting statistically significant enrichment in at least one of the foreground-region combinations; *p* values <0.05 are marked by asterisk. Sequence logos are shown on right side of the heatmap. The two foreground event sets and the three regions are indicated on the top of the heatmap. (C) Heatmap depicting Z-score of transformed TPM values of 9 candidate RBPs identified by the computational screening method.

KH RNA Binding Domain Containing, Signal Transduction Associated 3B, (*KHDRBS3*, *SLM2*, *T-STAR*) was one of the identified RBP splicing regulators with its RNA binding motif enriched in exon body region from negatively-correlated foreground event set and showed a temporal increase in expression during induced osteogenic differentiation (**Figure 3B, 3C**). Previous studies reported that *KHDRBS3* and another member from the STAR family, *KHDRBS1* (*SAM68*), are involved in splicing regulation in a variety of developmental and disease processes (Baralle and Giudice, 2017; Danilenko et al., 2017; Farini et al., 2020; Feracci et al., 2016; Traunmuller et al., 2016; Zhang et al., 2020). Examples of two significantly changing exon skipping events, unaccompanied by a difference in gene expression, are shown in **Figure S3**. During induced osteogenesis two potential *KHDRBS3* pre-mRNA targets, the transcription factor coregulator gene *AIRD4B* (*RBP1L1*, *RBBP1L1*) and the periostin (*POSTN)* gene were alternatively spliced but not differentially expressed. In both instances a KHDRBS3 RNA binding motif was present in the body of the regulated exon with the PSI value of the target exon being negatively correlated with *KHDRBS3* gene expression (**Figure S4, S3A, S3C**).

*ARID4B* and *ARID4A* (*RBP1*, *RBBP1*) are two homologous members of the AT-rich interaction domain (ARID) gene superfamily (Wilsker et al., 2005); they encode subunits of the SIN3 Transcription Regulator Family Member A (SIN3A)/HDAC (histone deacetylase) transcriptional corepressor complex which functions in various cellular processes including proliferation, differentiation and cell fate decision (Clark et al., 2015; Fleischer et al., 2003). As depicted in **Figure S3B**, the alternatively spliced cassette exon 16 in *ARID4B*, which encodes part of the Tudor-knot domain, is significantly more often skipped (PSI diminished) in day-12 compared to day-0 differentiating MSPC. Compared to the traditional Tudor domain, which is involved in protein-protein interaction (Lasko, 2010), the Tudor-knot contains crucial configurations needed for RNA binding activity (Shimojo et al., 2008). Therefore, partial disruption of the Tudor-knot during induced osteogenesis might result in changes in the assembly or stability of *ARID4B* involved supramolecular complexes. For example, in the mouse *Arid4a* is reported to be a *Runx2* coactivator that promotes osteoblastic differentiation (Monroe et al., 2010). Considering the fact that *ARID4A* and *ARID4B* can also physically interact with each other (Wu et al., 2013), the alteration of their protein-protein or protein-RNA interaction might squelch *ARID4A*’s function as a *RUNX2* coactivator.

Periostin is a secreted extracellular matrix protein that was originally identified in cells from the mesenchymal lineage in the skeleton (e.g., osteoblasts and osteoblast-derived cells). Periostin expression promotes bone anabolism partially though its ability to regulate osteoblast differentiation from mesenchymal progenitors (Bonnet et al., 2012); deletion of *Postn* gene impairs fracture consolidation in mice (Duchamp de Lageneste et al., 2018). As was the case with *ARID4B*, *POSTN* expression does not change significantly over the course of induced osteogenesis but exon skipping in *POSTN* pre-mRNA does, indicating that *POSTN* has the potential to contribute to osteogenic differentiation without a change in expression level. There are multiple isoforms of periostin reported, all differing in their C-terminal sequences (Litvin et al., 2004). This is consistent with our observation that the inclusion level of exon 18 which resides in the C-terminal region of *POSTN* decreased from 76% to 50% over the 12-day course of MSPC osteogenesis (**Figure S3D**).

### siRNA knockdown of KHDRBS3 reduces osteogenesis in vitro

As noted above, *KHDRBS3* was one of the nine RBPs that emerged from our computational screening pipeline (**Figure 3** and **Figure S4**). *KHDRBS3* showed a robust increase in expression over time during induced osteogenesis. This made *KHDRBS3* an ideal candidate for siRNA knockdown to test the biological significance of *KHDRBS3* during osteogenesis. siRNA knockdown of *KHDRBS3* relative to negative control siRNA was observed over seven days of induced osteogenic differentiation (**Figure 4**). As assessed by qPCR, a statistically significant knockdown of *KHDRBS3* to levels at or below initial levels of expression at all time points was achieved (**Figure 4A**). Diminished osteogenic differentiation was confirmed by significant reduction of the osteogenic maturation marker genes *RUNX2* (**Figure 4B**) and *BGLAP* (**Figure 4C**) in three and four time points, respectively. A significant reduction in functional readout of osteogenesis, alkaline phosphatase staining (**Figure 4D**) was also observed. These data indicate that KHDRBS3 expression normally supports osteogenesis perhaps via its action on downstream target genes (e.g., ARID4B and POSTN) to promote progression of MSPC maturation down the osteogenic pathway.

**Figure 4.**
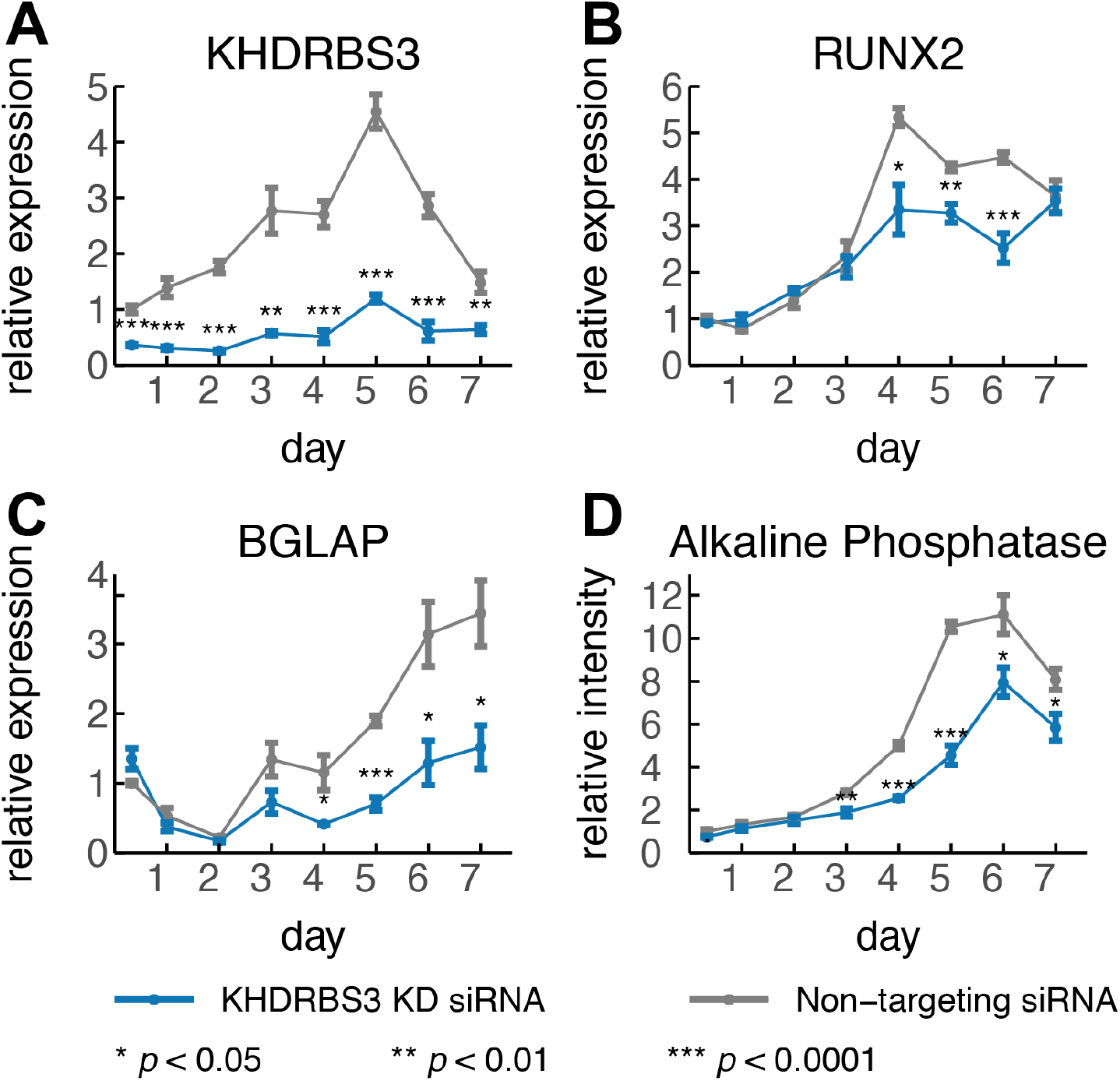
*KHDRBS3* knockdown reduced osteogenic differentiation of MSPC. *KHDRBS3*-specific siRNA knockdown (n=3) compared to non-targeting siRNA (n=3) at daily time-points upon exposure to osteogenic differentiation media was assayed for effects upon: (A) *KHDRBS3*, (B) *RUNX2*, and (C) *BGLAP* expression by qPCR. (D) plot of alkaline phosphatase activity staining with knockdown by *KHDRBS3*-specific siRNA compared to non-targeting siRNA. Error bars represent the MEAN±SEM (n=3). Asterisks indicate significance by two-tailed t-test.

## Discussion

While there have been some studies of AS events that occur during tissue-specific adipogenesis (Aprile et al., 2018; Huot et al., 2012; Li et al., 2014; Vernia et al., 2016), there is much less known about AS during the differentiation of mesenchymal progenitors down the osteogenic pathway in the human bone marrow niche. Studies of AS during osteogenesis have focused primarily on exon skipping in *RUNX2*, the master regulator gene for osteogenesis (Makita et al., 2008). Skipping exon 5 and/or 7 in *RUNX2* pre-mRNA produces isoforms of RUNX2 incapable of DNA binding and downstream transactivation of genes required for normal bone formation, including osterix (*OSX*), *OCN* (osteocalcin), *OPN* (osteopontin) and *COL1A1*. Some of these genes can also be regulated by AS. For example, *COL1A1* encodes alpha-1 type I collagen, which is the most plentiful collagen in bone. Aberrant AS of the *COL1A1* gives rise to a form of the human skeletal disease osteogenesis imperfecta (Johnson et al., 2000; Wang et al., 1996; Xia et al., 2008).

In this study, we generated a 12-day time-course RNA-seq dataset from primary cultures of MSPCs harvested from a human femur after they were induced to differentiate to bone-forming osteoblasts *in vitro*. This dataset not only permits a comprehensive examination of stepwise changes for both gene expression and AS, but also opens the opportunity to examine for the first time in a completely unbiased mode the potential regulatory role of RBP-directed changes in AS in MSPC differentiation. We found a high degree of similarity between the temporal patterns of overall gene expression, RBP gene expression and exon skipping (**Figure 1D**), suggesting that these events are mechanistically linked. We also observed that: 1) genes with the greatest variance in expression were significantly enriched for splicing-related gene ontology terms (**Figure S1B, S1C**); and 2) a large proportion of the differentially spliced genes encode transcription factors (**Supplementary Table 1**).

By combining temporal correlation of exon skipping and RBP expression with RBP binding site enrichment in the vicinity of regulated exons (**Figure 3**), we present a computational approach to identify key RBPs that drive AS changes in osteogenic differentiation. Considering the fact that AS is often regulated by binding of *trans*-acting RBPs to *cis*-acting RNA elements in a position-dependent manner (Fu and Ares, 2014; Park et al., 2018; Yee et al., 2019), it is important to understand how the region-specific RBP binding influences nearby AS events. Instead of using the whole set of differential AS events, for each candidate RBP two sets of AS events were determined based on positive or negative temporal correlation with RBP expression in osteogenic differentiation. RBP motif enrichment was assessed in three regions (upstream intron, exon body, downstream intron) as a proxy for region-specific RBP association. It should be noted that this computational strategy is generic and can be applied to any time-course RNA-seq dataset and to any type of AS patterns to elucidate RBP regulation of AS.

In this work, nine RBPs were identified as potential key splicing regulators in the process of MSPC to osteoblast differentiation. Among those was *KHDRBS3* that exhibited increased expression during osteogenesis (**Figure 3C**) and has been found to control cell fate in development or disease (Baralle and Giudice, 2017; Traunmuller et al., 2016; Zhang et al., 2020). As depicted in **Figure 4**, successful siRNA knockdown of *KHDRBS3* resulted in a commensurate decrease in functional readouts of osteogenesis, including alkaline phosphatase-directed bone formation. This result indicated that: 1) an increase in KHDRBS3 expression influences normal MSPC osteogenesis; and 2) our computational strategy was successful in identifying a key splicing regulator in the dynamic setting of MSPC-to-osteoblast differentiation. Further evidence is needed to prove that the function of KHDRBS3 on osteogenesis results from its regulation on splicing.

## METHODS

### MSPC culture

Primary cultures of MSPCs were obtained from PromoCell (C-12974, Heidelberg, Germany). Cells were characterized by the vendor according to criteria proposed by the International Society for Cellular Therapy (Dominici et al., 2006). Lot 402Z027 (47, male, Caucasian) was used in RNA-seq study. Lot 429Z013.1 (56, male, Caucasian) was used in siRNA knockdown study. MSPCs were initially cultured in recommended growth media (PromoCell, C28009) and differentiated in MSPC osteogenic differentiation medium (PromoCell, C-28013) on plates coated with human fibronectin (PromoCell, C-43060). Media was changed every two or three days. Samples for osteogenic differentiation RNA-seq data were obtained at day 0, 2, 4, 6, 8, and 12; and daily samples from day 0 to 7 for siRNA knockdown.

### Cell staining for biomarkers of osteogenic differentiation

Alkaline phosphatase staining reagent (5-Bromo-4-chloro-3-indolyl phosphate/Nitro blue tetrazolium) was prepared from BCIP/NBT tablet (Sigma B-5655, St. Louis, MO) in 10 ml water and incubated on cell monolayers after PBS wash for 10 minutes. BCIP/NBT reagent was removed by washing with PBS-Tween 0.05% followed by a PBS wash. Alkaline phosphatase staining was quantified by spectrophotometry at 620 nm. Alizarin Red S (ARS; Sigma A-5533) was prepared at 2% in water and adjusted to pH 4.1 and filtered before usage. Cells were fixed with 10% buffered formalin (Fisher) and washed with water prior to addition of 2% ARS, pH 4.1 for 20 minutes. Excess ARS stain was washed from cells by water four times. Staining was quantified by spectrophotometry at 405 nm.

### siRNA knockdown

*KHDRBS3* SMART pool On-Target Plus siRNA (Dharmacon) and On-Target Plus non-targeting pool siRNA (Dharmacon) was used as negative control for experiments. Transfection of 7000 cells per well (96-well plates) was conducted with X-treme GENE siRNA transfection reagent (Sigma) at 160 ng siRNA to 1 ul regent ratio. After eight hours, siRNA and transfection reagent containing media was removed and replaced with MSPC osteogenic differentiation medium (day 0). Media was changed every two or three days for a total of seven days. RNA was isolated from 96-well plates using RNeasy 96 (Qiagen). For qPCR gene expression analysis, cDNA was synthesized by SuperScript IV reverse transcriptase (Thermofisher) and qPCR performed with TaqMan Fast Advanced Master Mix (Thermofisher) with eukaryotic 18S rRNA endogenous control probe/primer (ThermoFisher) and gene specific probe/primers: *RUNX2* (Hs01047973_m1), *BGLAP* (Hs01587814_g1), and *KHDRBS3* (Hs00938827_m1). Staining was performed as described as above.

### RNA isolation and sequencing library preparation

For RNA-sequencing (RNA-seq), RNA was extracted from 24-well plate MSPC cultures at 0, 2, 4 6, 8 and 12 days of induced osteogenesis with Trizol (ThermoFisher) and purified with Direct-zol RNA microprep columns (Zymo Research). Three biological replicates were isolated at each time point. RNA-seq libraries were prepared with TruSeq Stranded mRNA Library Kit (lllumina) after which RNA was assessed for quality by Tape Station (Agilent) and quantified by Qubit (ThermoFisher). RNA-seq libraries were pooled, quantified by Qubit 3.0, diluted accordingly and committed to Illumina Paired End 101 base sequencing at the UCLA Broad Stem Cell Research Center High Throughput Sequencing Facility.

### RNA-seq read alignment

High-quality raw sequencing reads were obtained and assigned to a corresponding sample by demultiplexing with a maximum of 1 mismatch allowed in the barcode sequence (barcode sequence length 7). Alignment was done using *Hisat2* (v2.0.3-beta) (Kim et al., 2019) with default parameters and a pre-built index for reference plus transcripts based on genome assembly GRCh37 (hg19) annotation (grch37_tran, ftp://ftp.ccb.jhu.edu/pub/infphilo/hisat2/data/grch37_tran.tar.gz).

### Gene expression quantification and differential gene expression analysis

Gene expression/transcript abundance were measured in both raw counts and TPM (Transcripts Per Million) using the alignment tool kallisto (v0.43.1) (Bray et al., 2016). Ensemble v75 GRCh37 (hg19) cDNA annotation was used as the guiding reference for kallisto. Transcript-level estimates from kallisto were summarized into gene expression matrices by tximport (v1.6.0, R package) (Soneson et al., 2015) for downstream gene-level analysis. Differential expression analysis was conducted with the count-based tool DeSeq2 (v1.18.1, R package) (Love et al., 2014). Technical replicates were collapsed, and lowly expressed genes (TPM <= 5 in all samples) were filtered out before performing differential expression analysis. For each comparison, genes with an absolute log2 fold change > log2(1.5) and an FDR (false discovery rate)-adjusted p-value < 0.01 were assumed to be differentially expressed genes. The differentially expressed gene list for the entire osteogenic differentiation pathway was defined as genes differentially expressed in all comparisons between time point day 0 and other time points (day 2, 4, 6, 8, 12).

### Alternative splicing analysis to identify significantly changing foreground events and background events

AS events were detected and quantified by rMATS-turbo (Shen et al., 2014), with Ensemble v75 GRCH37 (hg19) GTF annotation. Exon inclusion levels, measured as PSI values, were calculated by junction reads (reads spanning the splicing junctions) normalized by effective junction length. AS events with low junction read support (≤ 10 average junction reads, ≤ 10 total inclusion junction reads or ≤10 total skipping junction reads over all 18 samples), or with extreme PSI value ranges (PSI ≤ 0.05 or ≥ 0.95 in all 18 samples) were excluded from downstream analysis. Differential exon skipping analysis was then performed using rMATS-turbo (with default parameter -c 0.0001) for five pair-wise comparisons between time point day 0 and other available time points (day 2, 4, 6, 8, 12). Exon skipping events for each comparison were considered differential if they meet the following criteria 1) >10 average junction reads (inclusion and skipping junction reads) in both groups; 2) does not have extreme PSI values (PSI ≤ 0.05 or PSI ≥ 0.95 for all 6 samples in the comparison); 3) FDR < 0.01; and 4) absolute change in PSI (|∆PSI|) > 0.05. The significant event set for the whole osteogenic differentiation pathway was composed of events that were identified to be differentially spliced in at least 3 of the 5 comparisons. The background event set for the whole osteogenic differentiation pathway was defined as events with no significant change during MSPC osteogenic differentiation which meet the following cutoffs in all 5 comparisons: 1) > 10 average junction reads in both groups; 2) does not have extreme PSI values (PSI ≤ 0.05 or PSI ≥ 0.95 for all 6 samples in the comparison; and 3) FDR > 0.5.

### Principle component analysis (PCA)

A total of 129 RBPs (including many well-characterized splicing factors) were curated from two different sources (Dominguez et al., 2018; Ray et al., 2013) and included in PCA analysis of RBP expression. For total gene or RBP expression, a pseudo-count of 1 was added to each TPM value before log2 transformation to avoid arithmetic error and large negative values. PCA was then performed after removing genes/RBPs/exon skipping events with no variance among samples. Samples were projected to their PC1-PC2 space by PCA score. LOESS (locally estimated scatterplot smoothing) regression lines with 95% confidence intervals were added to the PC1-PC2 plot using R package ggplot2 (v3.1.0).

### Gene set enrichment analysis (GSEA)

GSEA (v3.0) software (Subramanian et al., 2005) was utilized on pre-ranked gene lists based on absolute values of principle component loadings from PCA. Ranked lists from total gene expression PCA included only the top 10,000 genes; for ranked lists from exon skipping, duplicated genes with lower rank were removed. All gene ontology gene sets (c5, v7.0, https://www.gsea-msigdb.org/gsea/msigdb/download_file.jsp?filePath=/msigdb/release/7.0/c5.all.v7.0.symbols.gmt) were used as gene sets database with 1000 permutations to calculate the enrichment score and p-values. Top gene ontology terms from the GSEA analysis served as input for REViGO webserver (http://revigo.irb.hr/) to account for the semantic similarities and dispersibilities of gene sets. Representative gene ontology terms were visualized in semantic similarity-based scatter plots after removing redundant ones.

### Hierarchical clustering of time course datasets and heatmaps

Hierarchical clustering was performed on Z-score transformed TPMs of differentially expressed RBPs (from a total of 1542 RBPs) (Gerstberger et al., 2014) or PSI values of differentially spliced exons detected in the whole osteogenesis pathway as described above (hclust function in R package stats, v3.4.4).

### Protein family domain analysis

Pfam domain scanning was conducted to search for potential functional domains affected by exon skipping events. Preprocessed Pfam annotation data, which maps HMM predicted high-quality Pfam-A domains to UCSC hg19 coordinates, were downloaded from the UCSC hg19 annotation database (http://hgdownload.soe.ucsc.edu/goldenPath/hg19/database/ucscGenePfam.txt.gz). Sequences of target exons or frameshifted downstream exons from differential exon skipping events were extracted and then scanned against the Pfam annotation data using bedtools (v2.25.0) (Quinlan and Hall, 2010).

### RBP candidate screening method

189 RBP binding motifs with position weight matrix information for 129 RBPs (including many well-characterized splicing factors) were curated from two different sources and screened in this analysis. This includes 78 6-mer motifs for 78 RBPs from RNA Bind-n-Seq (RBNS) (Dominguez et al., 2018) and 111 7-mer motifs for 82 RBPs from RNAcompete (Ray et al., 2013).

Significant/background event lists and differentially expressed RBPs were identified as described before (see AS analysis section and gene expression analysis section). For each motif of the differentially expressed RBPs, significant exon skipping events were further assigned to two foreground event sets. They were composed of events (n > 50) whose PSI value was positively or negatively correlated (R^2^ > 0.5, rlm function in R package MASS, v7.3-49) with differential RBP gene expression across different time points.

To identify region-specific RBP regulatory patterns for exon skipping events, we evaluated three regions around the alternatively spliced exons: 1) 300 nt of intronic sequence upstream of the target exon; 2) the exon body sequences; and 3) 300 nt of intronic sequence downstream the target exon. Scores for each motif were calculated by sliding window scanning of the position weight matrix at each possible binding position. Region-specific motif occurrence was then determined by comparing the calculated motif scores with a threshold score (80% of the maximum PWM score). If there was any position with a calculated motif score ≥ the threshold score for a particular exon skipping event, then the motif occurrence was marked as “True” for this event in the corresponding region; otherwise it was marked “False”.

To determine whether a motif occurred in a specific region more often in foreground event sets than in the background event set, we used a one-tailed Fisher’s exact test to test the null hypothesis that the number of events with motif occurrence at a specific region was not different between the foreground (either positive or negative set) and the background event set. If an RBP motif was of significantly enriched occurrence (p <0.05) in any region for any foreground event set, it was considered a key RBP for exon skipping regulation in the MSPC osteogenic differentiation process.

---

**Supplementary Figure 1.**
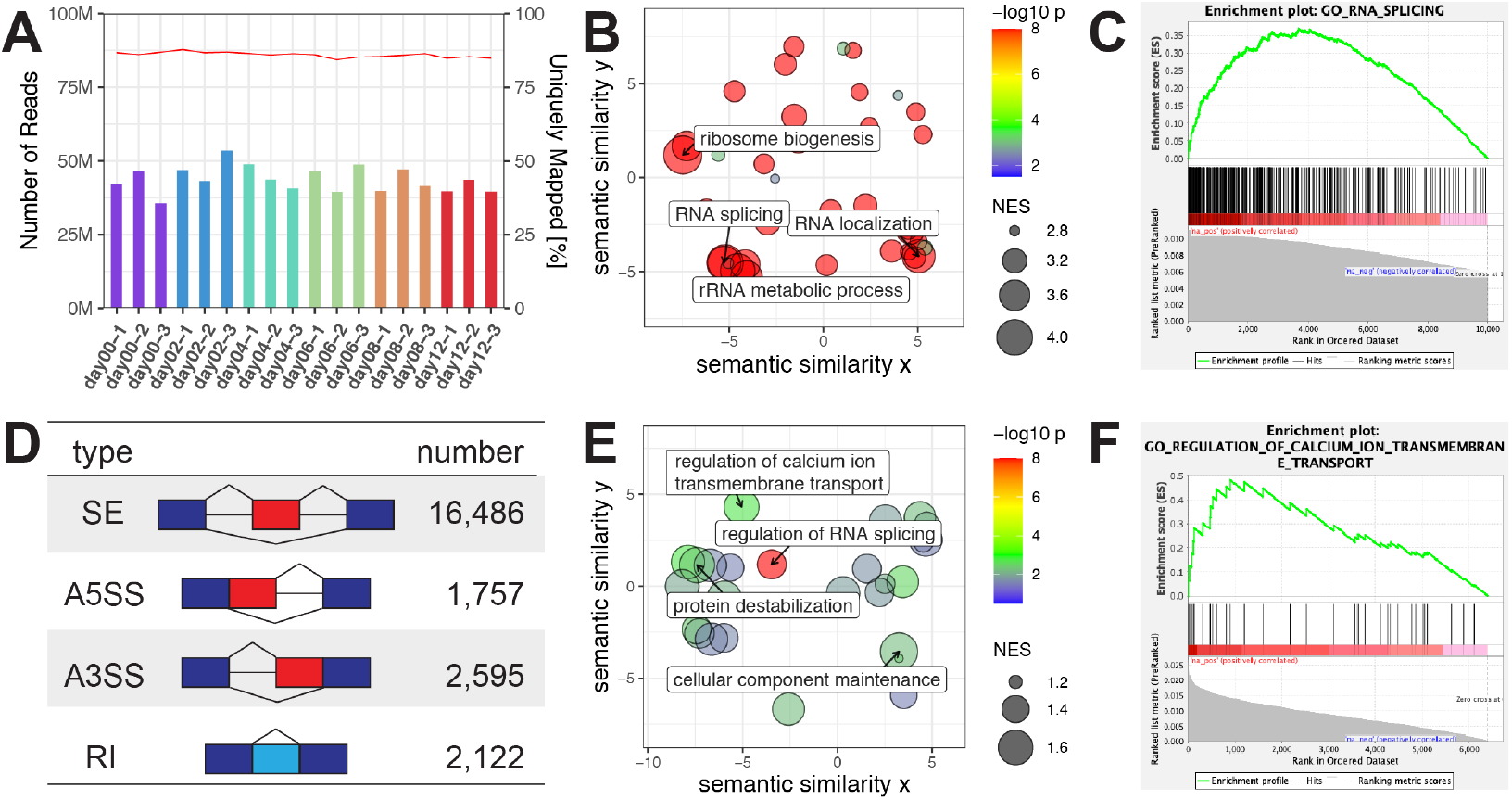
Transcriptome-wide analysis of osteogenic differentiation identifies interplay of gene expression, splicing and bone development related biological processes. (A) Summary of read depth and mapping statistics from RNA-Seq dataset. (B) REVIGO scatter plot depicting gene ontology terms enriched among genes with high PC1 loading from total gene expression PCA (see left panel, **Figure 1D**). Gene ontology enrichment was evaluated by gene set enrichment analysis (GSEA). GSEA calculated enrichment scores and p-values are indicated by size of circles and color scale, respectively; x- and y-axis represent the semantic similarities between terms. Representative gene ontology terms were labeled. (C) Representative GSEA enrichment plot (GO_RNA_SPLICING) from panel A. (D) Summary table of AS events detected by rMATS-turbo after filtering by read coverage and PSI value range. SE, exon skipping; A5SS, alternative 5’ splice sites; A3SS, alternative 3’ splice sites; RI, intron retention. (E) REVIGO scatter plot depicting GO terms enriched among genes with high PC2 loading from exon skipping PCA (see right panel, **Figure 1D**). (F) Representative GSEA enrichment plot (GO_REGULATION_OF_CALCIUM_ION_TRANSMEMBRANE_TRANSPORT) from panel E.

**Supplementary Figure 2.**
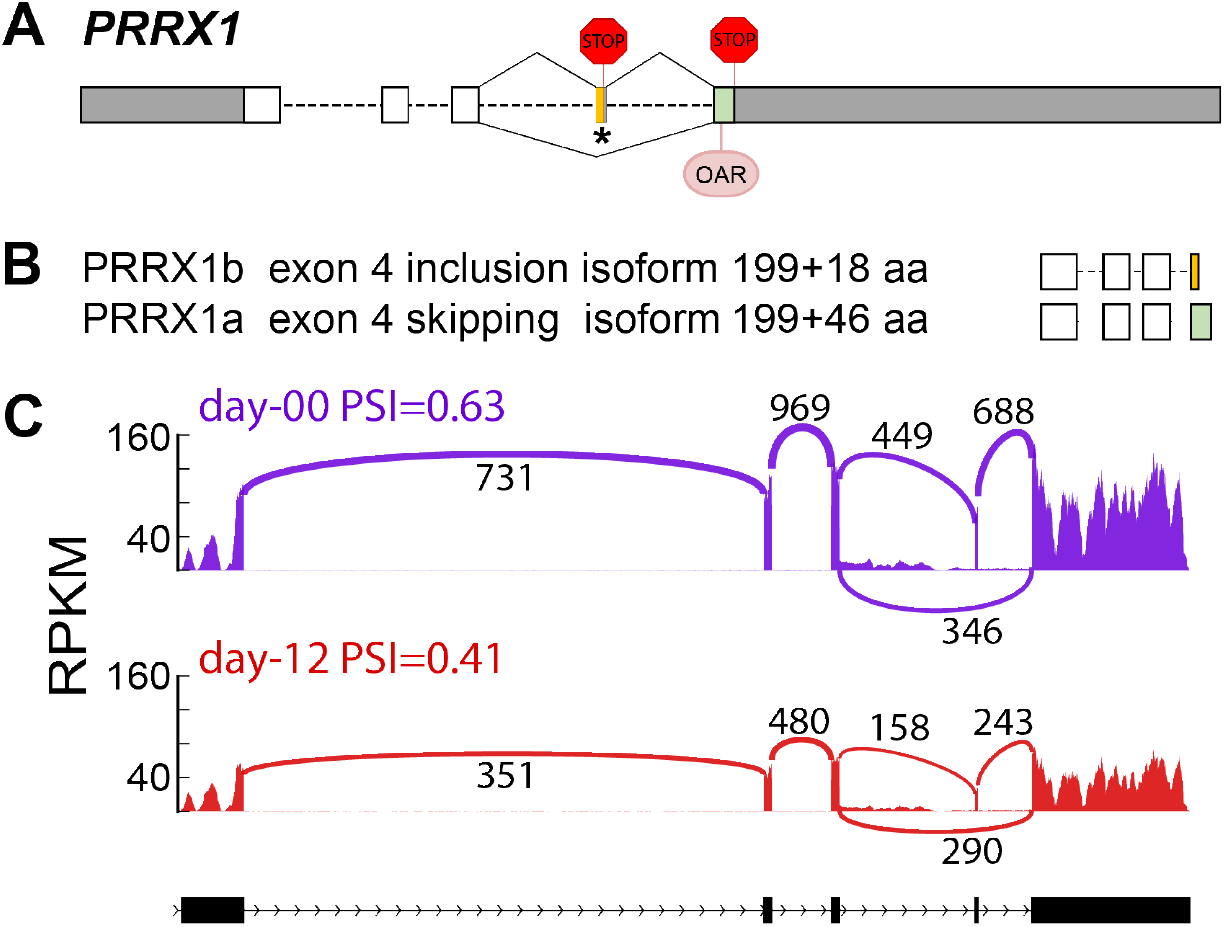
Exon 4 inclusion/exclusion of *PRRX1* results in an isoform switch. (A) Gene structure representation of the *PRRX1* gene. Bars and dashed lines represent exons and introns, respectively. Untranslated regions (UTR) are denoted in grey. Two stop codons residing in exon 4 and 5 are depicted as STOP signs. The OAR (otp, aristaless, and rax) domain is located in the green region in exon 5. (B) Protein structure representation of the two *PRRX1* isoforms. The top PRRX1b isoform has exon 4 incorporated and the bottom PRRX1a isoform has exon 4 excluded. The stop codon in exon 4 of the inclusion isoform renders exon 5 and OAR domain untranslated. (C) Sashimi plot showing AS changes of the *PRRX1* gene on day 0 (upper panel) and day12 (lower panel). The black bars and dashed lines on the bottom represent exons and introns, respectively. Solid peaks represent RPKM of reads mapped to each region. Arches represent splice junctions and the numbers represent number of reads mapped to each splice junction.

**Supplementary Figure 3.**
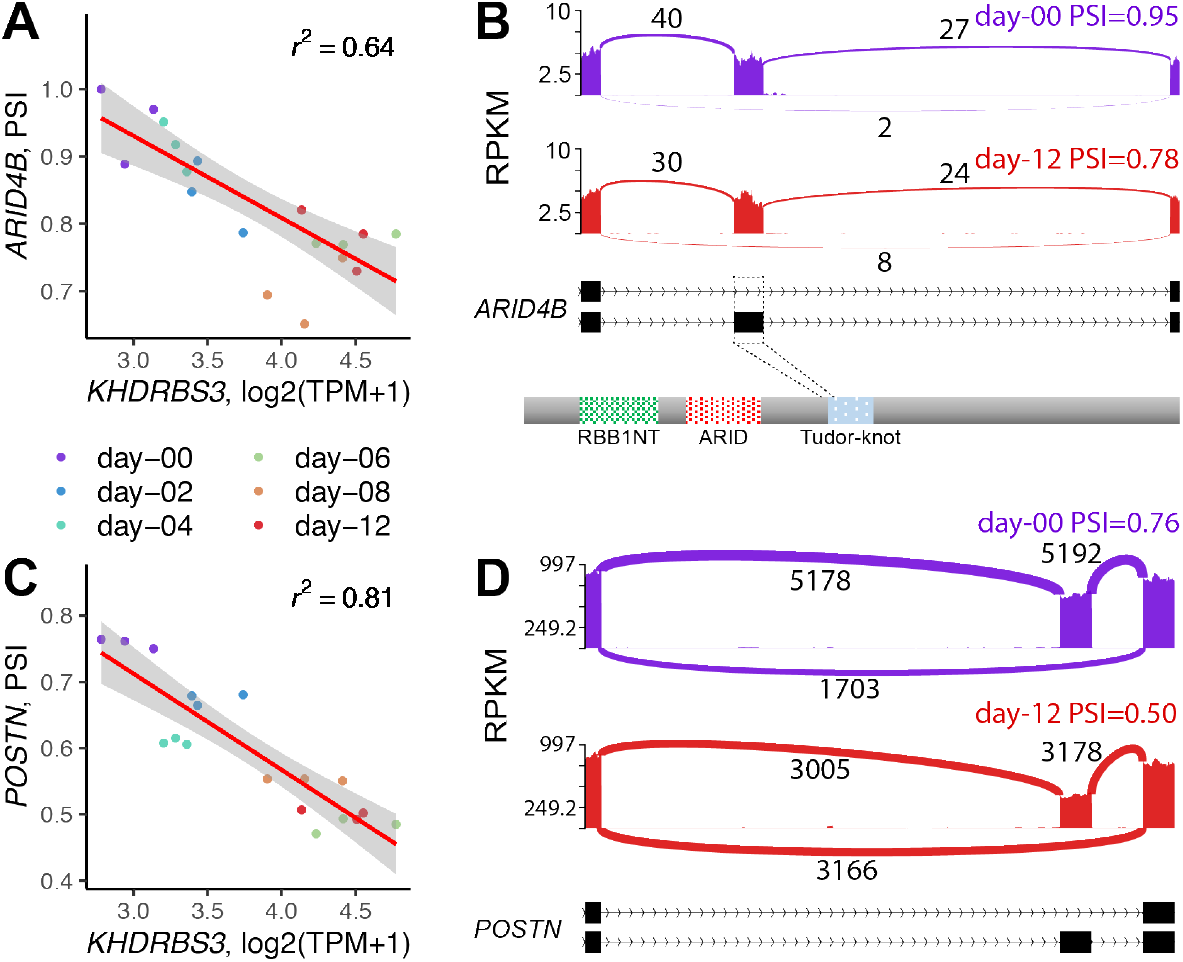
Examples of exon skipping events in putative targets of KHDRBS3. (A, C) Correlation between exon skipping (PSI) and *KHDRBS3* gene expression changes over 12 days of induced osteogenesis in MSPC. Genes harboring the significantly changing exon skipping event are indicated on the y-axis. Linear regression lines (red) and confidence intervals (grey) are shown for correlated *KHDRBS3* expression and *ARID4B* PSI (A) and POSTN PSI (C) during induced osteogenesis and r^2^ values of the correlation are shown in the upper right corner. (B) Sashimi plot of an exon skipping event in the transcription factor gene *ARID4B*. The black bars and dashed lines on the bottom represent exons and introns, respectively. Solid peaks represent RPKM mapped to each region. Arches represent splice junctions and the numbers of reads mapped to each splice junction. On the bottom is the depiction of protein family (Pfam) domains in ARID4B protein. (D) Sashimi plot of the exon skipping event in gene *POSTN*. The target exon 18 is located in the Cterminal sequence of POSTN, where extensive AS changes occur.

**Supplementary Figure 4.**
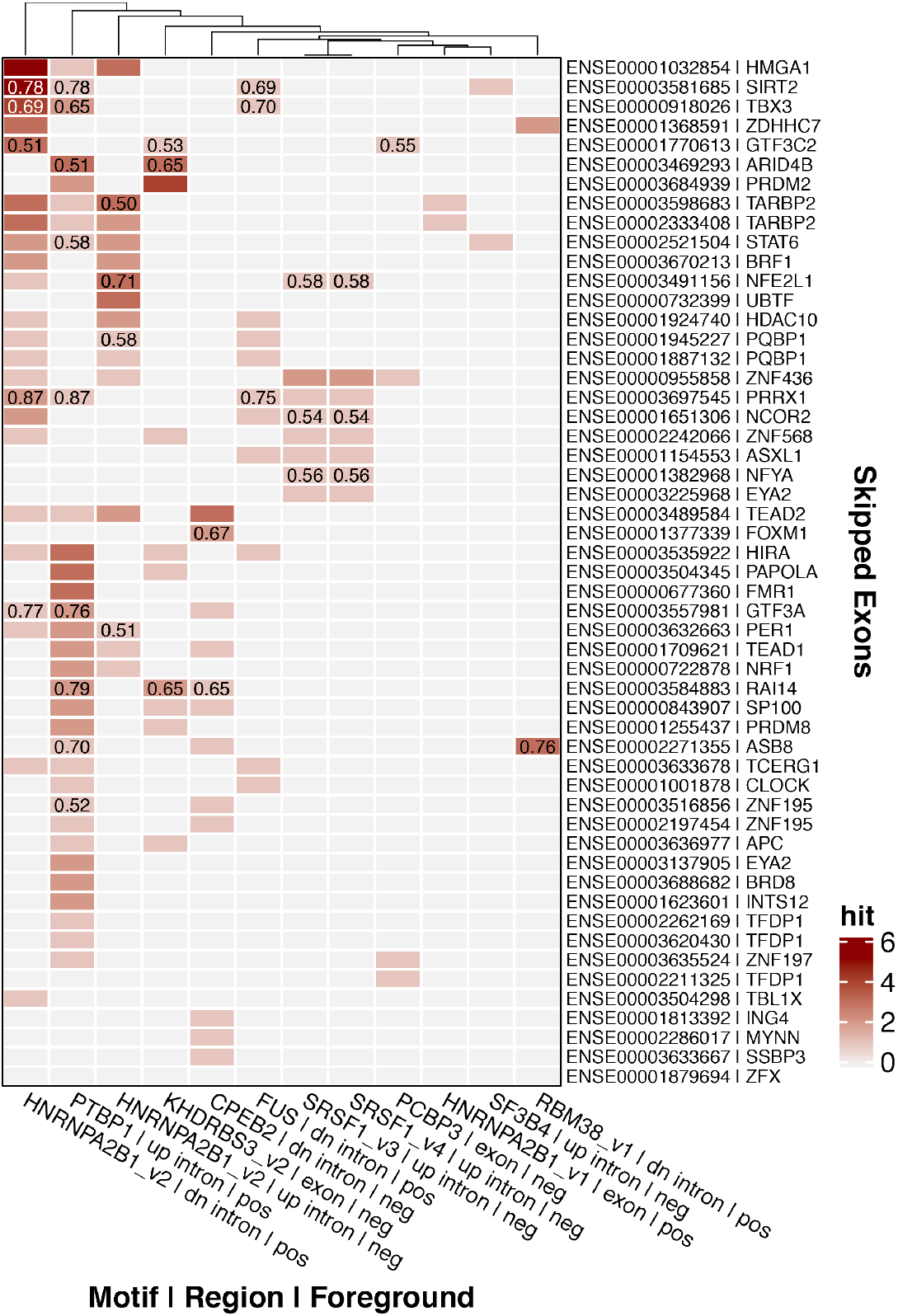
Heatmap of candidate RBP motif hits in exon skipping events occurring in a transcription factor. Candidate RBP motifs (from **Figure 3B**) are indicated along the x-axis; additional information includes the region of significant RBP binding motif enrichment (up intron=upstream intron, exon=exon body, dn intron=downstream intron) and direction of correlation (pos=positive or neg=negative). Posted along the y-axis are the gene symbols of the transcription factor and exon identification number in that transcription factor where the differential exon skipping event was detected. The colors of heatmap represent motif occurrence in the designated region. r^2^ values > 0.5 for the correlation between RBP gene expression and PSI value of the exon skipping event are indicated numerically in the appropriate boxes.

**Supplementary Table 1.**
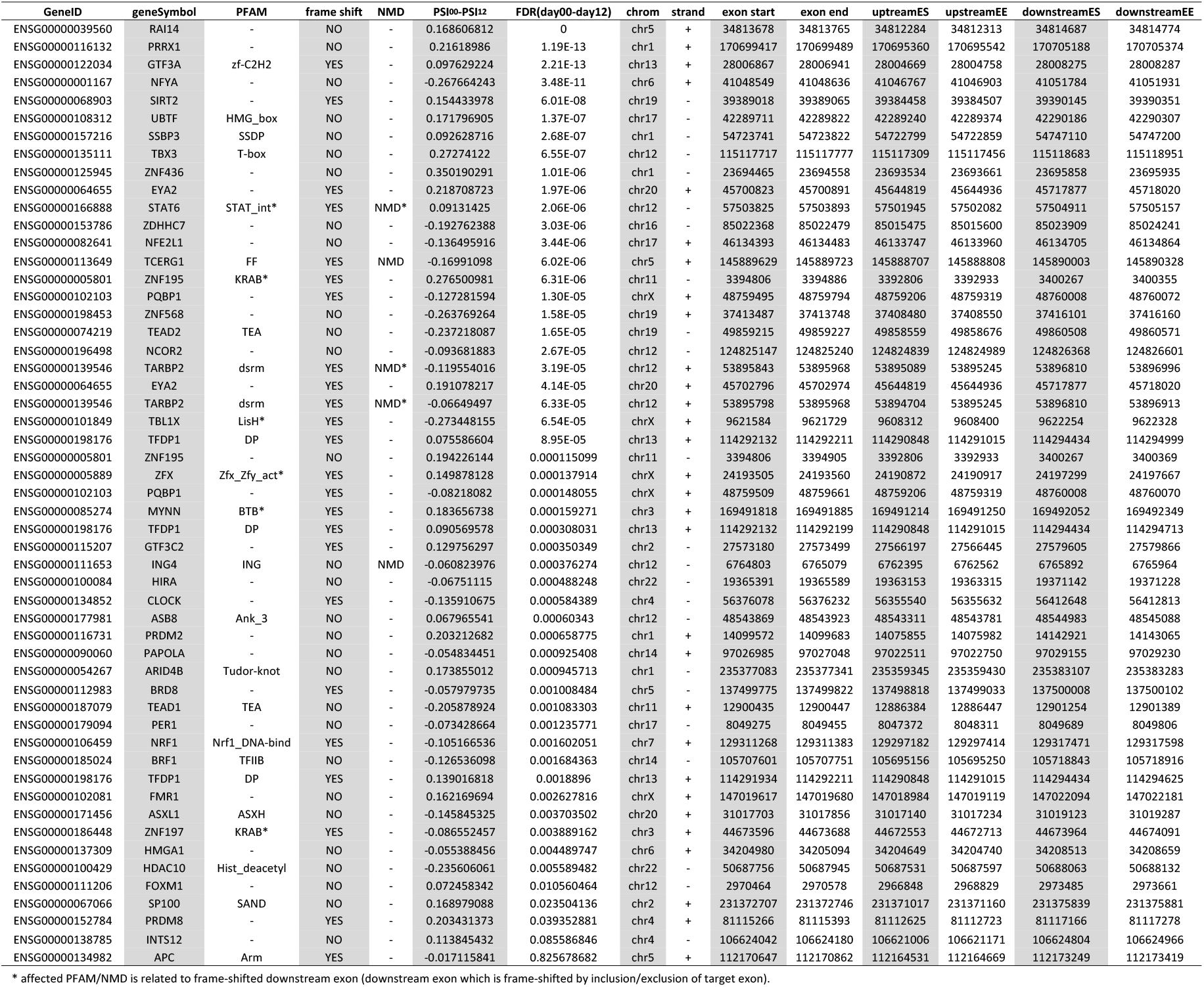
Transcription factors with exon skipping events are often affected by frame shift and/or disruption of functional domains. Shown are transcription factors with significant exon skipping changes during osteogenic differentiation. Column 3 reports the protein family domain (Pfam) affected by inclusion/exclusion of target exon, due to the presence/absence of the domain, encoded by the target exon or the downstream exon (*) which is frame-shifted by incorporation/exclusion of target exon. Column 4 indicates whether the target exon incorporation results in a frameshift. Column 5 indicates whether the target exon or the frameshifted downstream exon (*) is involved in nonsense mediated decay.

